# Assessing Selectivity in the Basal Ganglia: The “Gearbox” Hypothesis

**DOI:** 10.1101/197129

**Authors:** Zafeirios Fountas, Murray Shanahan

## Abstract

Despite experimental evidence, the literature so far contains no systematic attempt to address the impact of cortical oscillations on the ability of the basal ganglia (BG) to select. In this study, we employed a state-of-the-art spiking neural model of the BG circuitry and investigated the effectiveness of this circuitry as an action selection device. We found that cortical frequency, phase, dopamine and the examined time scale, all have a very important impact on this process. Our simulations resulted in a canonical profile of selectivity, termed selectivity portraits, which suggests that the cortex is the structure that determines whether selection will be performed in the BG and what strategy will be utilized. Some frequency ranges promote the exploitation of highly salient actions, others promote the exploration of alternative options, while the remaining frequencies halt the selection process. Based on this behaviour, we propose that the BG circuitry can be viewed as the “gearbox” of action selection. Coalitions of rhythmic cortical areas are able to switch between a repertoire of available BG modes which, in turn, change the course of information flow within the cortico-BG-thalamo-cortical loop. Dopamine, akin to “control pedals”, either stops or initiates a decision, while cortical frequencies, as a “gear lever”, determine whether a decision can be triggered and what type of decision this will be. Finally, we identified a selection cycle with a period of around 200ms, which was used to assess the biological plausibility of the popular cognitive architectures.

**Author summary:** Our brains are continuously called to select the most appropriate action between alternative competing choices. A plethora of evidence and theoretical work indicates that a fundamental brain region called the basal ganglia might be the locus where this competition occurs. But how is the winning choice determined each time? Using a detailed computational model, based on neurophysiological properties of this region, we suggest that, whereas the basal ganglia might indeed contain the circuitry of action selection, the cerebral cortex is, in fact, the brain region that dictates this process. Similarly to a gearbox in a car, the basal ganglia provide modes for the exploitation of the safest option (forward gears), exploration of alternative options (reverse gear) and a neutral state, in case that the selection process needs to be halted. Our results further indicate that the instructions for mode-switching are relayed to the basal ganglia through specific low frequencies of oscillations within cortical areas. Finally, we provide estimations for the frequency ranges that can be used to activate each selectivity mode, as well as the duration of the selection process under various conditions.

## Introduction

The physical location of the basal ganglia (BG), as well as their broad bidirectional connectivity with major cortical areas, the limbic system and the thalamus, place the this brain structure in a key position to modulate the flow of information between the cortex and the body. Despite the great diversity of inputs and outputs, the human BG consist of the same repeating internal circuitry [1] which is also largely retained in most vertebrate species [2,3]. This strictly topographic organization on different scales suggests that through this structure, some common modulatory operations are applied to functionally different channels of information flow.

In the microscopic scale, the BG circuitry can be broken down into a massive number o parallel loops (or channels) which, as suggested, represent different competing information signals or “action requests” [4,5]. According to this popular theory, the BG circuit is able to process those requests and finally select the most salient (or urgent) potential action, via the direct BG pathway, while providing inhibition to the rest competing channels via the indirect pathway [6,7].

An increasing amount of neurophysiological evidence implicates the BG to selection of voluntary motor actions and provides indirect verification of this hypothesis [8]. [9] showed that the excess activation of the direct BG pathway in freely behaving mice, via stimulation of MSN_*D*1_ neurons in the striatum, increases movement, while the stimulation of the indirect pathway made the same animals to freeze. In addition, although both pathways are required for healthy action selection and were found to contribute equally to the initiation of actions in [10], the indirect pathway is suppressed during the execution of actions or action sequences [11], presumably because any behavioural conflicts have already been resolved during movement [8].

From another standpoint, low-frequency brain oscillations have been widely implicated in both the function of the BG [12] and the process of decision making [13–17]. Oscillations in the cortex mediate the processing of new information [18], the dynamic formation of neural ensembles representing different actions and the suppression of other task-irrelevant regions [19,20]. The are also found to encode uncertainty and influence the exploration-exploitation trade-off [21]. In addition, there is a substantial number of studies focusing on low-frequency oscillations in the BG, as changes of this activity are connected with a number of disorders such as Parkinson’s or Huntington’s disease.

But are these phenomena related? Evidence suggests that oscillations in some certain bands in the striatum and the subthalamic nucleus (STN), the input structures of the BG, are driven by cortical regions [22–25]. Taking this into account, in previous work [26] we explored the impact of cortical rhythmic activity on the BG function and we found that the former can completely shape which areas of the BG circuit are active. Yet, the connection between the BG, cortical oscillations and decision making still remains relatively unexplored.

In this work we attempt to narrow this gap by investigating whether cortical oscillations could influence the ability of the BG to act as a selection device. To achieve this, we initially defined a number of metrics that enable the assessment of the effectiveness of possible selection mechanisms. Using the biologically plausible neural model of the BG circuitry defined in [26], we then carried out an analysis of the relationship between cortical frequencies, dopamine concentration and BG selectivity.

We found that the frequency and phase difference between oscillatory cortical areas, the level of dopamine in the system and the examined time scale, all have a very important impact to the ability of our model to select. Our simulations resulted in a canonical profile of selectivity in the BG, which we termed selectivity portraits, that can be largely maintained in simplified versions of the model.

Using these portraits, we show that although the BG circuit can robustly and sequentially perform selection tasks, the strongly-active cortical areas instruct the mode of this selection via their oscillatory activity. Some frequency ranges promote the exploitation of actions of which the outcome is known, others promote the exploration of new actions with high uncertainty, while others simply deactivate the selection mechanism. Finally, we identified a selection cycle with a period of around 200 ms, which was used to assess the biological plausibility of the most popular architectures in cognitive science.

Our results can be replicated with simpler versions of the neural model used, agree well with experimental observations, provide new justifications and insights into oscillatory phenomena related to decision making and reaffirm the role of the BG as the selection centre of the brain.

## Materials and Methods

### The basal ganglia model

#### The full model

The predominant tool used in this study is a state-of-the-art, large-scale spiking neuron model of the complete motor BG circuitry, presented in detail in [26]. This model integrates fine-tuned models of phenomenological (Izhikevich) spiking neurons that correspond to different sub-types of cells within the BG nuclei, electrical and conductance-based chemical synapses that include short-term plasticity and neuromodulation, as well as anatomically-derived striatal connectivity.

In particular, this model comprises 10 neural populations that correspond to the four major nuclei of the biological BG and form the canonical circuit described in the Introduction. These include the striatum and the STN, the two inputs of the BG, the external part of the globus pallidus (GPe), as well as the substantia nigra pars reticulata (SNr), one of the two output structures of the BG. Furthermore, the effect of the pars compacta part of the substantia nigra (SNc) is realized through the concentration of the neurotransmitter dopamine (DA) in the different parts of the network (green colour in Fig. 1). The network is devided in three microscopic channels, which are mutually inhibited and used to represent different action requests throughout this study.

**Fig 1.**
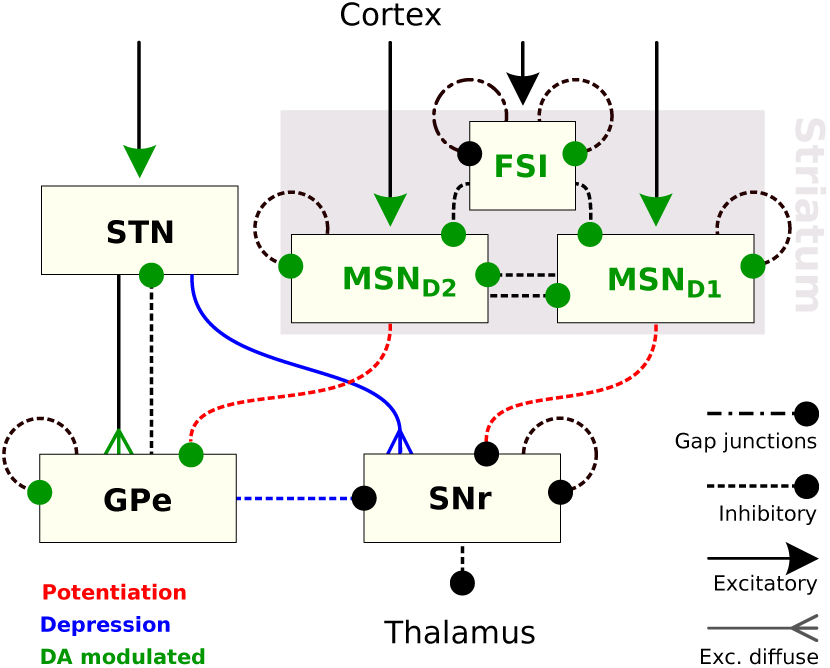
The network architecture of the current BG model.

The internal structure of the striatum has been modelled using three different groups that correspond to its three major neural populations. The first two groups, which comprise the 99% of the striatum, correspond to the two categories of medium spiny-projection neurons (MSNs), divided based on the dominant type of their dopamine receptors. Dopamine is known to enhance activity in the first group (MSN_*D*_1 neurons) and depress the second (MSN_*D*_2 neurons). The remaining 1% of the striatum is occupied by fast-spiking gabaergic interneurons (FSIs) that are affected by both types of dopamine receptors and are highly interconnected with both electrical and GABAergic synapses. Finally, the STN and GPe comprise three sub-populations each, that correspond to the three predominant types found in the literature to have distinctive dynamical and electrophysiological patterns.

The number and ratio of neurons in each group, is taken from anatomical studies and result in a total of 9586 neurons that form the BG network. The probability for a connection between any two neurons of this network depends on the source and target nuclei and it was either infered from anatomical studies or taken from previous computational studies. Finallly, an optimization method was used to approximate neural and network parameters based on empirical findings [27].

#### A reduced version of the model

As a consequence of its detailed architecture, the neural model in [26] contains a rather high number of parameters that might influence its behaviour and the resulting measurements of selectivity. In order to narrow down this space and establish the most important BG features for selectivity we defined a second, simplified version of this neural model with significantly less differences between nuclei. The behaviour of this simplified model is compared against the full version in [26] in the Results section of this study, where the homogeneity and robustness of our results is determined. The architecture of the simplified model is shown in Fig. 2.

**Fig 2.**
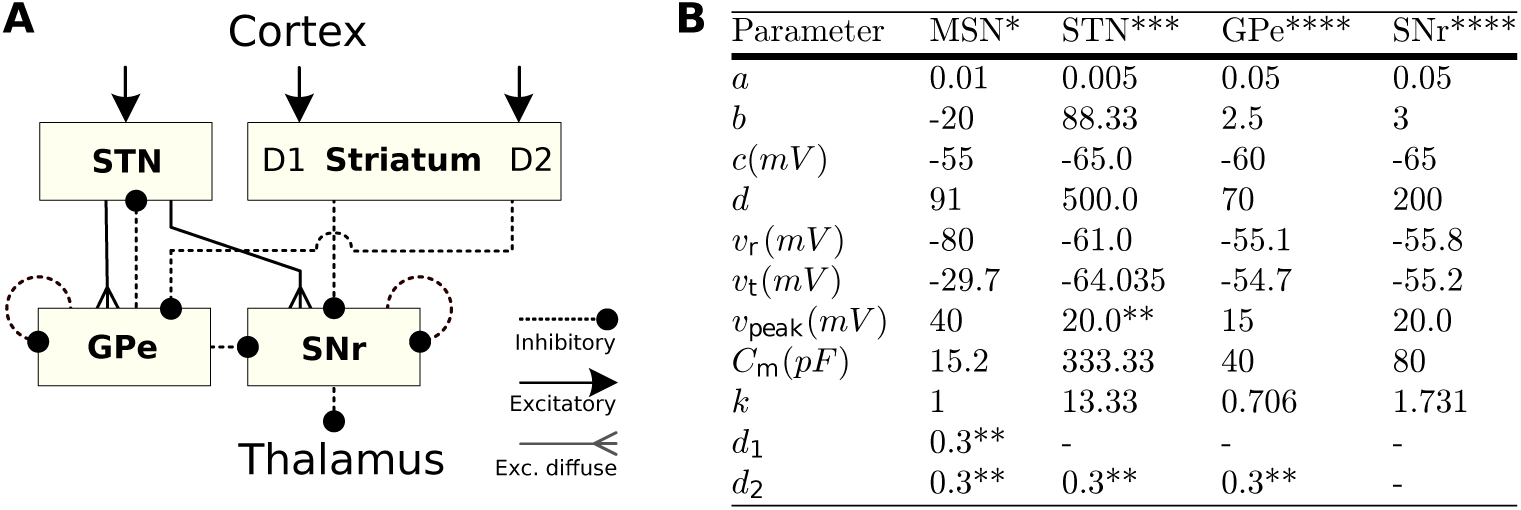
Simplified version of the BG model. * Parameters taken from [28] ** Parameters taken from [4] *** Parameters derived from [29] **** Parameters derived from [30]

Here, the striatum is modelled using only 600 D1-like and 600 D2-like MSNs with the FSIs and gap junctions being neglected due to their small number. The rest neuron groups consist of the STN, GPe and SNr, which were modeled using a single parameter, set as well as a fixed number of 150 neurons for each group. The values for all neuron parameters can be found in the table of Fig. 2B. The synapses between neurons in this model do not exhibit short-term plasticity. They include AMPA, NMDA and GABA types and they are governed by conductance-based equations. They have fixed reversal potentials E equal to 0, 0 and −80 for each neurotransmitter respectively,*τ_AMPA_* = 2,*τ_NMDA_* = 100 and *τ_gaba_* = 3, as well as a maximum conductance *g* = 1*nS* for all connections in the system. In addition, no optimization was conducted to fit the firing rates of the model to the corresponding biological nuclei, as it was the case with the original model in [26]. Instead, the probability for each neuron of a source nucleus to be connected to a neuron in the target nucleus was always set to 0.25. Finally, the cortical input towards the three microscopic channels of this model remained the same as in the case of the full version of thr BG model, in order to enable a more direct comparison.

### Metrics

#### Selectivity

The view of the BG as the action selection device implies that their performance on this aspect could be evaluated based on measurable criteria, such as signal distinction. The further suggestion that the salience of an action is encoded in the local level of activity in the striatum and STN, which is directly affected by cortical input, can serve as the basis of this evaluation. [31] defined selectivity in the BG as the ability of a neural mechanism to robustly distinguish competing signals. Although this definition is sufficient, the main focus of this study was confined to the difference between transient and steady-state effects, and it produced metrics that can not be applied in a more general case, such as the BG model of the current study.

Our aim here is to create a metric that is aligned with the features of our model but it also remains general enough to be used in other studies. The first step of this attempt is to find a method to measure the *distinctiveness* of a single selected channel. This can be defined as the ability of a channel to receive distinctively less inhibition than any other channel or, more specifically, the degree to which the following conditions are fulfilled: (a) The firing rate of the selected channel in the level of the SNr is close to zero, which is required in order to revoke inhibition in the thalamus, and (b) no other channel is far below tonic levels. These two conditions can be written as

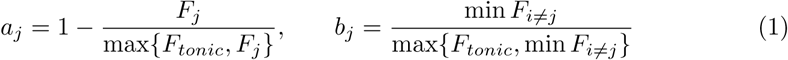

where *j* is the examined channel, *F_i_* is the SNr firing rate of a channel *i* and *F_tonic_* is the tonic firing rate of the SNr (~ 25 spikes/sec). Since both denominators in (1) are upper-bounded by the value of the corresponding numerator, the product *D̅_j_* = *a_j_b_j_* will always take values between [0,1] and reflects the requested measure. The special case of *F_j_* = min *F_i≠j_* = *F_tonic_*/2 results in 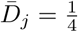 and represents the baseline below which the channel *j* is indistinguishable. To normalize *D̅_j_*, so the baseline lies in 0, the final *distinctiveness D_j_* of a channel *j* is given as

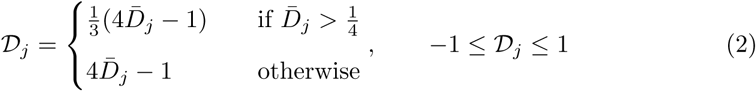

A graphical illustration of the above can be found in Fig. 3A. Using this metric we can now measure a number of properties of the BG selection mechanism. First, the *effectiveness* of the BG in selecting the most salient cortical input can be defined as

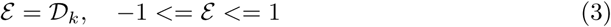

**Fig 3.**
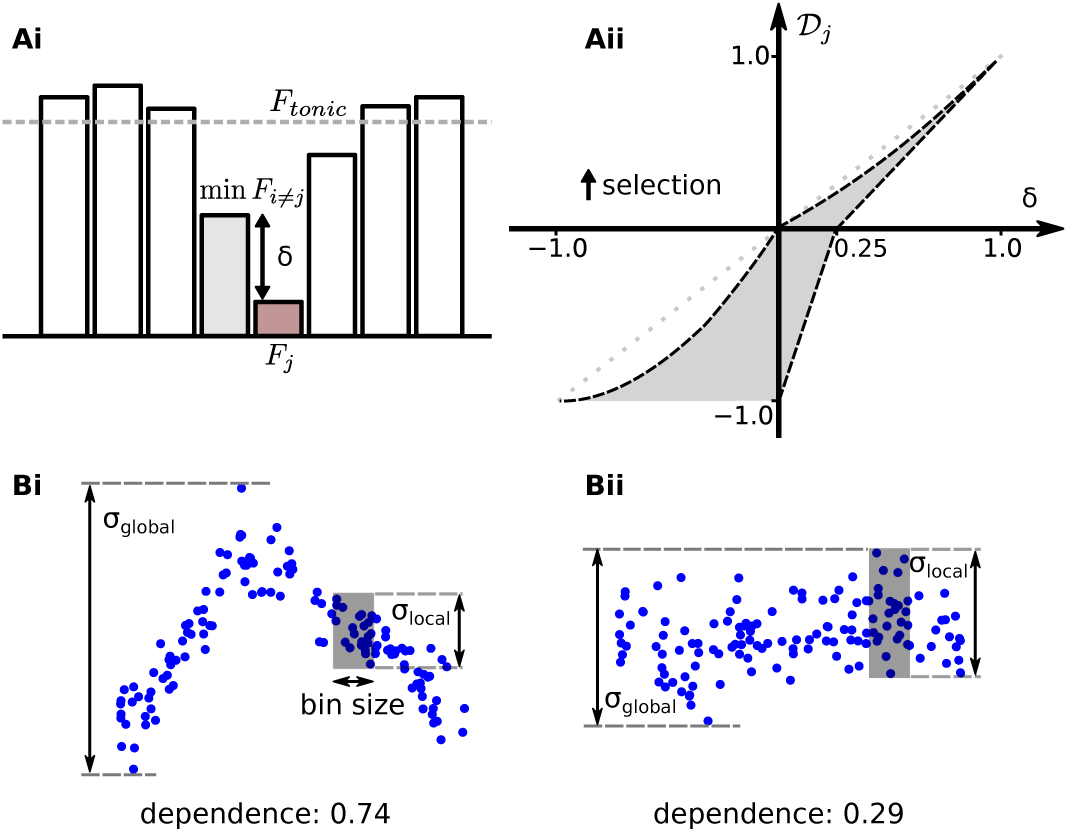
Metrics for distinctiveness and dependence. **Ai:** A multi-channel example of SNr firing rates used to illustrate the concept of distinctiveness of the channel *j*. **Aii:** Space of possible values for *D_j_* for any difference *δ* between *j* and the least-inhibited alternative channel. Note that the line *D_j_* = *δ* is an upper bound to the possible values of *D_j_* in this space. **B:** Example calculation of dependence.

where *k* is the index of the most salient channel, i.e. the channel with the highest firing rate at the level of the cortex.

Furthermore, the degree of *selectivity* of the BG reflects to their ability to select *any* channel regardless of its salience and can be defined as

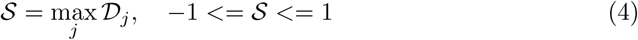

Finally, one more useful property that can be measured using *D_j_* is to what extent the BG is selecting, or exploring, alternative actions. This is given as

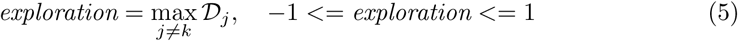

and is defined as the level of exploratory behaviour of the BG mechanism, or simply *exploration*. To compare this metric with terminology commonly found in the literature the value of effectiveness in the BG can be considered here as the level of *exploitativeness*, since the high salience of the leading microscopic channel arises from a previously learnt behaviour. Hence, selectivity can be then thought of as the union of exploration and exploitation.

To conclude, *D_j_* can be used to measure various features of a neural-based action selection mechanism with minimal adjustments. The only requirements are first, a local measurement of the instantaneous firing rate in the output area of a neural structure, and second, a prior knowledge of the average tonic firing rate in the same area. In case that the latter cannot be obtained, the difference *δ* between the selected channel and the least-inhibited neighbouring area (Fig. 3Aii) provides a good approximation of distinctiveness, especially when 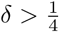, and thus it can be used instead.

#### Transient versus steady-state

An event processed by a selection mechanism can have both a transient and a steady-state effect on a dynamical system such as the brain. Our BG model exhibited rich transient phenomena during the first 500*ms* after the injection of a stimulus, as well as a different post-transient steady state that was maintained indefinitely. To distinguish between these two modes, the *transient* distinctiveness of a salient channel is defined as the maximum degree by which this channel received less inhibition than any other neighbouring channel for a fixed short interval, after the generation of the salient signal. That is

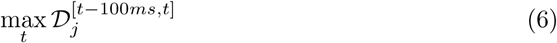

where *T* + 100*ms* < *t* < *T* + 500*ms* and *T* denotes the point on time that the stimulation was applied. The *steady-state* distinctiveness, on the other hand, can be measured taking into account the post-transient stable firing rates in the level of the SNr.

#### Dependence

Selectivity can be affected by various parameters of the model or the current stimulus. Some of these parameters can play a decisive role in determining the model’s performance. The degree by which the BG selectivity depends on the value of a single parameter of the model can be measured by comparing the local versus the global variation of the resulting *D_j_* for a number of simulation runs with random initial conditions.

For instance, in case that selectivity is highly dependent on the value of a parameter, a significantly large sample of randomized simulation runs will result in diverse local mean values and small local standard deviations, compared to the standard deviation of the complete sample. An illustration of this concept is shown in Fig. 3Bi-ii.

This metric was used here to examine the effect of the phase offset *φ* between two oscillatory cortical inputs. In this case, local areas can be found by dividing the range of possible values for this parameter *R* = [0, 2*π*) into a number of bins *R_i_* = {*x*/*x* ∈ [*a*, *a* + *dx*), 2*π* · *a* = *i* · *dx*}, where *dx* is the length of each bin. Additionally, if *σ_i_* is the standard deviation of selectivity values within the bin *R_i_* and *σ_global_* the global standard deviation in *R*, the *dependence* of the BG selectivity to *φ* can be defined as

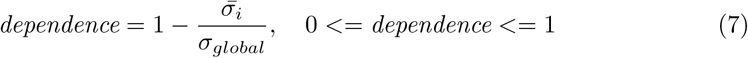

For the analysis of this study we have used 30 local areas (bins) to calculate dependence, a number which was found to provide adequate and robust results.

## Results

### Selectivity portraits

Initially, we conducted a series of simulations where the BG circuitry was called to resolve a conflict between two salient potential actions. To simulate this scenario, the BG model received strong cortical input in two out of their three microscopic channels, governed by 1000 inhomogeneous Poisson processes each, described in [26], and background noise of 3 spikes/sec in the third channel. These two strong inputs were oscillatory, with a single fixed frequency *f* = *f*_1_ = *f*_2_, but different amplitudes *A*_1_ < *A*_2_. Since the firing rate of the cortical ensembles that generate these inputs represents the salience of each action, the second cortical input was always considered the most salient one or, in other words, “the right choice”.

To investigate the relation between dopamine, cortical oscillations and the efficiency of the BG as a selection mechanism, we varied the frequency *f* of the two cortical ensembles, the phase offset *φ* between them, and the level of dopamine *d* = *d*_1_ = *d*_2_ in the system. An overview of the resulting BG behaviour can be seen in Fig. 4.

**Fig 4.**
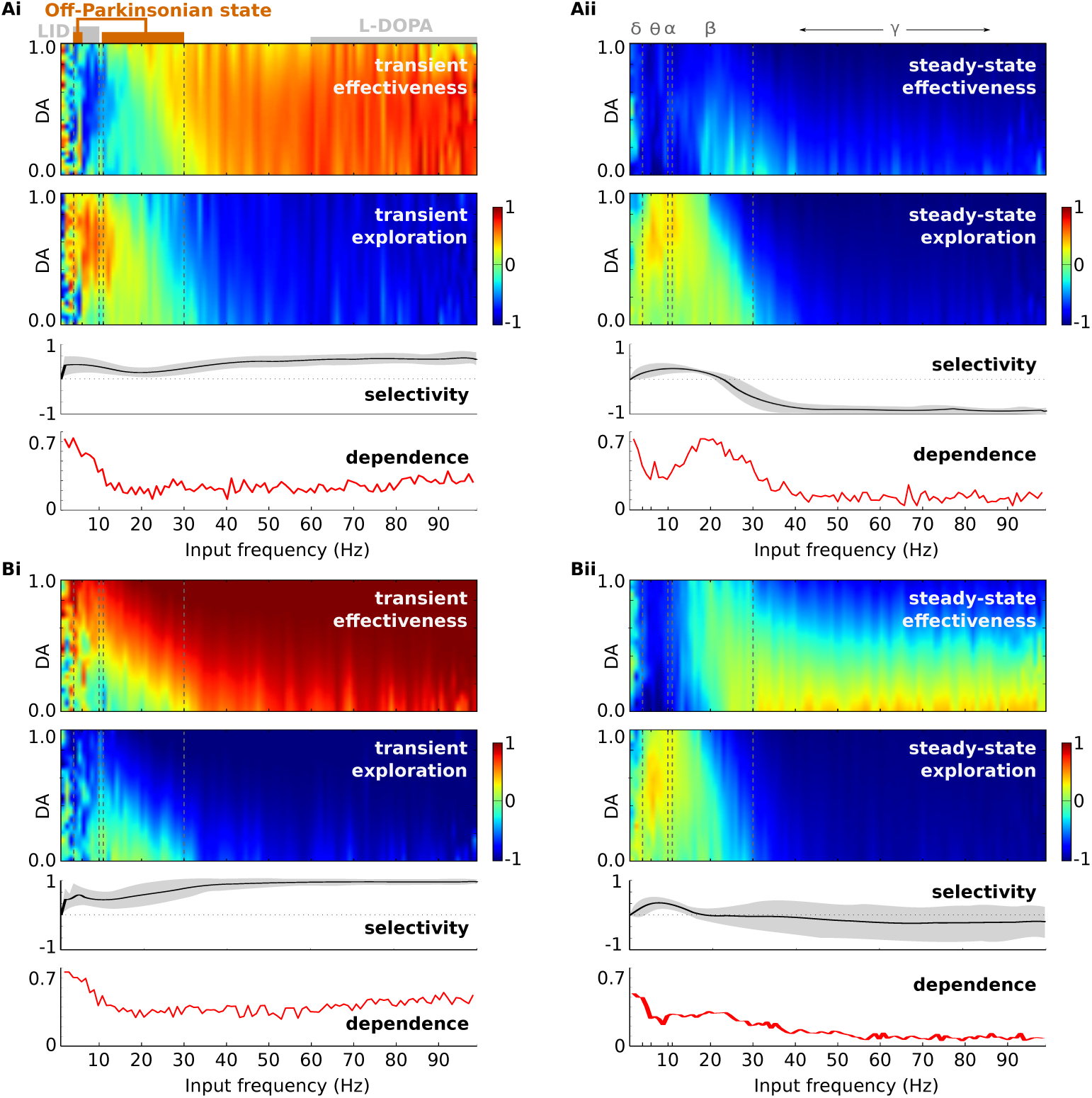
Selectivity portraits of the BG model. Effectiveness (scatter plot up), exploration (scatter plot down), selectivity (black curve) and dependence on the phase offset *φ* (red curve) when two inputs oscillate with amplitudes *A*_1_ = 7.5, *A*_2_ = 10 spikes/sec in (**A**) and *A*_1_ = 5, *A*_2_ = 10 spikes/sec in (**B**), in order to simulate strong and weak competition respectively. Cortical input to the third channel has a fixed baseline firing rate of 3 spikes/sec. Effectiveness is calculated for each combination of dopamine levels *d* and input frequencies *f*. The colour bars represent the mean of a sample of 200 runs (for each point) with random *φ* ∈ [0, 2*π*). Selectivity curves represent the mean (black line) and standard deviation gray area for all *φ* and *d*, across frequency spectrum. Dependence was calculated for *d* = 0.3.

The coloured scatter plots of this figure illustrate the tendency of the model to select the most salient signal (effectiveness), or the alternative, less-salient signal (exploration) for any possible combination of dopamine and cortical frequency. Finally, the plots right below indicate the ability of the system to select any signal, as well as the degree by which *φ* affects these measurements, across the same frequency spectrum. Since these figures can expose the critical conditions that affect the selection mechanism under examination, we termed them “*selectivity portraits*” of the model.

In the next paragraphs, we present a number of observations which were largely based on this figure, and we outline the most important testable predictions that emerged, regarding the function of the BG in the brain.

#### The combination of dopamine concentration and cortical frequency defines BG effectiveness and exploration

Fig. 4 clearly indicates that both the frequency of the two oscillatory inputs as well as the level of dopamine in the system play a crucial role in the ability of the BG to select. The responses of the model for various values of these two parameters revealed three main areas of interest in the frequency spectrum with completely different behaviour. The first area includes low-frequency oscillations, with a borderline at *f* = 15Hz, the second area corresponds to beta oscillations (13 < *f* < 30Hz) and the third area includes all greater frequencies.

In all cases, dopamine exhibited distinct patterns with which it regulated effectiveness and exploration. These patterns were completely different during the initial transient phase as opposed to the final steady-state BG response, while they were further modified depending on whether the competition was strong or week. As a result, the model generated four unique selectivity portraits when it dealt with each of the above cases.

More specifically, we found that, in our model, dopamine concentration affects selectivity only in particular frequency ranges, where its role is to either trigger or block the selection process. Notably, decisions triggered by dopamine promoted exploration over exploitation in the majority of the simulated scenarios. An exception is the case of a strong initial lead in the salience of the one of the competing channels before the level of the BG, showed in Fig. 4Bi. As the dominance of this channel is clear, an increased level of dopamine triggers the selection of this instead of the alternative choice. However, even if the most salient channel has already been selected transiently, this selection can be maintained over time only if dopamine decreases (Fig. 4Bii).

From the perspective of the cortical behaviour, low-frequency oscillations also promoted the selection of the least salient channel. This was achieved via the level of dopamine, which determined whether a selection will be made or delayed. With this type of input, the BG model became completely unable to maintain the first choice after an initial short transient.

On the other hand, beta oscillations minimized the influence of dopamine and brought the system in a neutral state, where both effectiveness and exploration are in the borderline value 0. Once more, this effect was halted in the presence of a strong difference between the two inputs.

Finally, gamma oscillations can clearly facilitate BG effectiveness. Transiently, selectivity maximized and the most salient channel was selected for any frequency, phase offset and dopamine level. In steady-state, gamma oscillations continued to support the same decision, but only if the channel remained highly salient and the level of dopamine dropped below tonic levels.

In order to ascertain the validity of these results and rule out the possibility that other stochastic parameters of the model had an important impact, we examined the variance of these measures when the examined parameters were fixed. Specifically, we ran 100 experiments where, each time, the level of dopamine, the input frequency and the phase offset *φ* were kept fixed to a random value within the biologically realistic limits but all other statistically defined entities in the model were randomised. These included the synaptic indexes, neural parameter perturbations and neuron types within a nucleus among others. This process was repeated 500 times giving in total 500 random points in the selectivity portraits that can be used for this analysis.

As a result, the three selectivity metrics presented Fig. 4 were almost identical between runs. A Shapiro-Wilk’s test [32,33] showed that the vast majority of these data points were approximately normally distributed, with an average p value *p* = 0.56 ± 0.36 that could not reject the null hypothesis of normality. The resulting standard deviations in each point were on average 0.114 ± 0.035 for effectiveness and 0.053 ± 0.032 for exploration.

The magnitude of this variation was very small compared to the differences in the selectivity process, and it was also comparable to the standard deviation of the normalized firing rates in the 3 channels of the SNr (0.072(±0.057) × 25 spikes/sec). Since these values are the only parameters of *D̅_j_*, our results indicate that there is no hidden correlation in the system, and the fluctuations of the standard deviation in selectivity plots of Fig. 4 were caused by dopamine and *φ*.

#### The BG can almost always select the most salient action transiently

During the simulations that produced the selectivity portraits, the BG model exhibited a significantly more aggressive selectivity transiently, at the first 500 ms after the presentation of the stimulus, as opposed to its steady-state behaviour. This is an expected range of reaction times in psychophysical choice tasks. It is consistent with oscillatory changes in the BG [34] and sensorimotor cortex [35] during animal decision making tasks, as well as choice reaction times in mental chronometry studies in humans [36-38]. However, the equation (6) that has been used to produce the selectivity portraits of the current model, does not fully address the dynamic changes of selectivity. A further comparison with experimental studies, such as the above, requires information regarding the onset and duration of the emerging transient peaks, as well as any rebound effects. The average response of our model for the four major examined frequency ranges is presented in Fig. 5.

**Fig 5.**
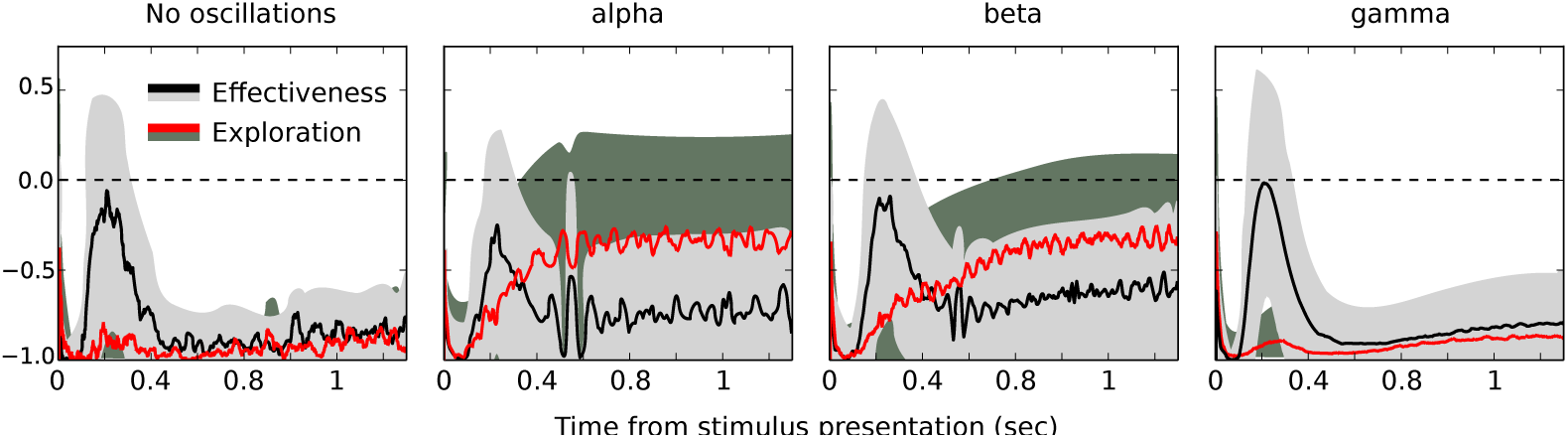
Transient changes in selectivity. Two competing cortical inputs oscillate with amplitudes 7.5 / 10 spikes/sec respectively. The two coloured curves represent mean value for any level of dopamine and offset *φ*, and the two coloured areas smoothed standard deviation.

As in the previous section, the large variations at some frequency ranges of this figure come from the different values of phase offset **φ** and the level of dopamine. For instance, beta frequencies cause positive effectiveness only when dopamine is greater than 0.8. Since successful selection cannot occur for *D_j_* < 0, we consider the BG as able to select only in scenarios where a significant portion of our experiments had a selectivity peak above this baseline.

Right after the presentation of the stimulus the BG model did not produce any selection response for a short period with fixed duration. Instead, the firing rate in all SNr channels was high, indicating an initial STOP phase. This phase had a very similar duration of 85 ± 67 ms on average, in all frequency ranges. Next, a transient increase in effectiveness that peaked at 133 ± 155 ms on average, accompanied the initial STOP phase. Although this increase had also a similar onset at all frequencies, its exact duration and the rebound activity varied significantly between the four frequency ranges (Average duration without oscillations: 81 ± 62 ms, in alpha oscillations: 42. ± 59 ms, beta: 28 ± 46 ms and gamma: 70 ± 63 ms). Hence, our results indicate that cortical frequency does not influence the reaction time of the BG, although different frequency ranges cause different types of reactions.

Furthermore, the model was not able to maintain effectiveness above the baseline after the first 500 ms. An exception to this rule was the case of alpha oscillations, where effectiveness had a second sharp rebound spike, with a surprisingly similar duration and onset among runs. Indeed, in some trials at these frequencies, selectivity was stronger in this second peak. This bimodal distribution of maximum selectivity between trials could reflect to a similar pattern in behavioural tasks. The latencies of the two peaks in our simulations are consistent with the bimodal distribution of reaction times in distinct cue choice tasks with rats [35]. However, the mechanism that caused this second selectivity peak is not yet fully understood, thus further investigation is required in order to establish its biological importance.

#### Cortical oscillations with low frequencies are required for selection change

The steady-state patterns of BG selectivity are also worth closer examination. Their function can be plausibly linked to a number of cognitive operations related to action selection. These include the ability of the BG to maintain a selection, for example during postural activities [34], to easily switch the current selection to an alternative cue, or the level of general alertness.

In Fig.4, it is shown that the most critical areas that affect effectiveness and exploration are mainly located in low frequencies while gamma oscillations have no discernible effect. In fact, Fig. 5 shows that gamma frequencies have virtually the same effect on selectivity as no oscillations.

To shed more light into the steady-state behaviour of the BG after the presentation of two competing stimuli, Fig.6 illustrates the firing rate of the BG output nucleus, the SNr, during that period and for the complete examined frequency spectrum.

**Fig 6.**
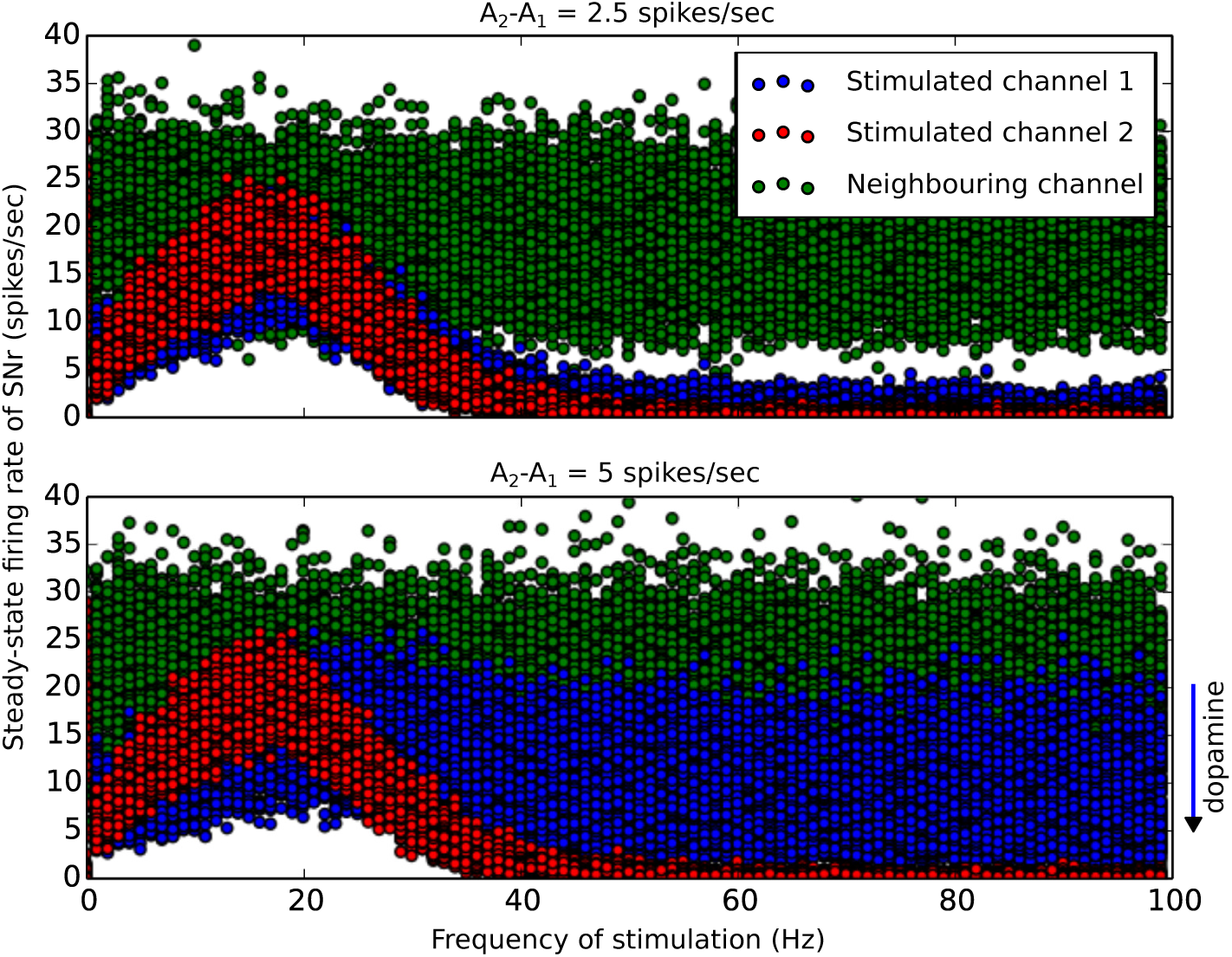
Inhibition of the SNr microscopic channels. **Up:** Firing rates of the three simulated microscopic channels in SNr, for various stimulation frequencies when two channels are stimulated with maximum amplitudes *A*_1_ = 7.5 and *A*_2_ = 10 spikes/sec respectively. **Down:** The same figure but for *A*_1_ =5 and *A*_2_ = 10 spikes/sec. Activity in channel 1 is reversely proportional to the level of dopamine in the system.

At low oscillations, and particularly at alpha frequencies, the firing rate of the selected microscopic channel was always close to the firing rate of tonic areas of this nucleus (25 spikes/sec). When the difference between the competing signals was low, this gave a clear advantage to the less salient channel which, under some conditions, could be directly selected. However, when the competition was less ambiguous, the advantage of the less salient channel diminished. In fact, during cortical oscillations at 20 Hz, the two salient channels were treated equally. They were both inhibited to around 50% of their default tonic state and, as a result, both channels remained ready for immediate deployment.

This specific beta frequency was of particular importance, since it manifested a critical state in the model. Higher cortical frequencies favoured the most salient channel and, on average, they significantly increased its distinctiveness, while low frequencies below 20 Hz had the exact opposite effect.

Finally, gamma oscillations also showed an interesting effect. In experiments with low ambiguity between competing channels and for low levels of dopamine, a selection of the highest salient channel could be maintained. However, under high ambiguity, or when dopamine increased, both channels remained inhibited, i.e. selected. This mechanism, which presumably allows information to flow via the cortico-BG-thalamic loop, might keep both information channels active until more evidence is accumulated. Cortical gamma synchronization has been widely associated with active information processing and feature binding [18,39,40]. Hence, the multi-selection mechanism we observed here might also contribute to these cognitive functions, by promoting integration of multiple information channels and thus allowing coalitions of neural ensembles to be formed.

#### Selectivity portraits are largely maintained in simplified versions of the BG model but not in the minimal model

The BG model that was developed in [26] and used in this study has an advanced degree of complexity. Although its behaviour is similar to its biological counterpart, the extent to which our results depend on specific modelling features or how robust they are for small perturbations requires further clarification. In an attempt to address these issues, we classified the individual features of the model according to their impact on selectivity portraits. To do this, we ran the same simulations shown in Fig. 4 but for each set of data points created, a single feature of the model was either disabled, or had its parameters randomized. When necessary, the optimization process for the connectivity of the model was repeated under these conditions, to bring the firing rates of the BG nuclei back to their biologically realistic ranges. The result of this classification is illustrated on Fig. 7.

**Fig 7.**
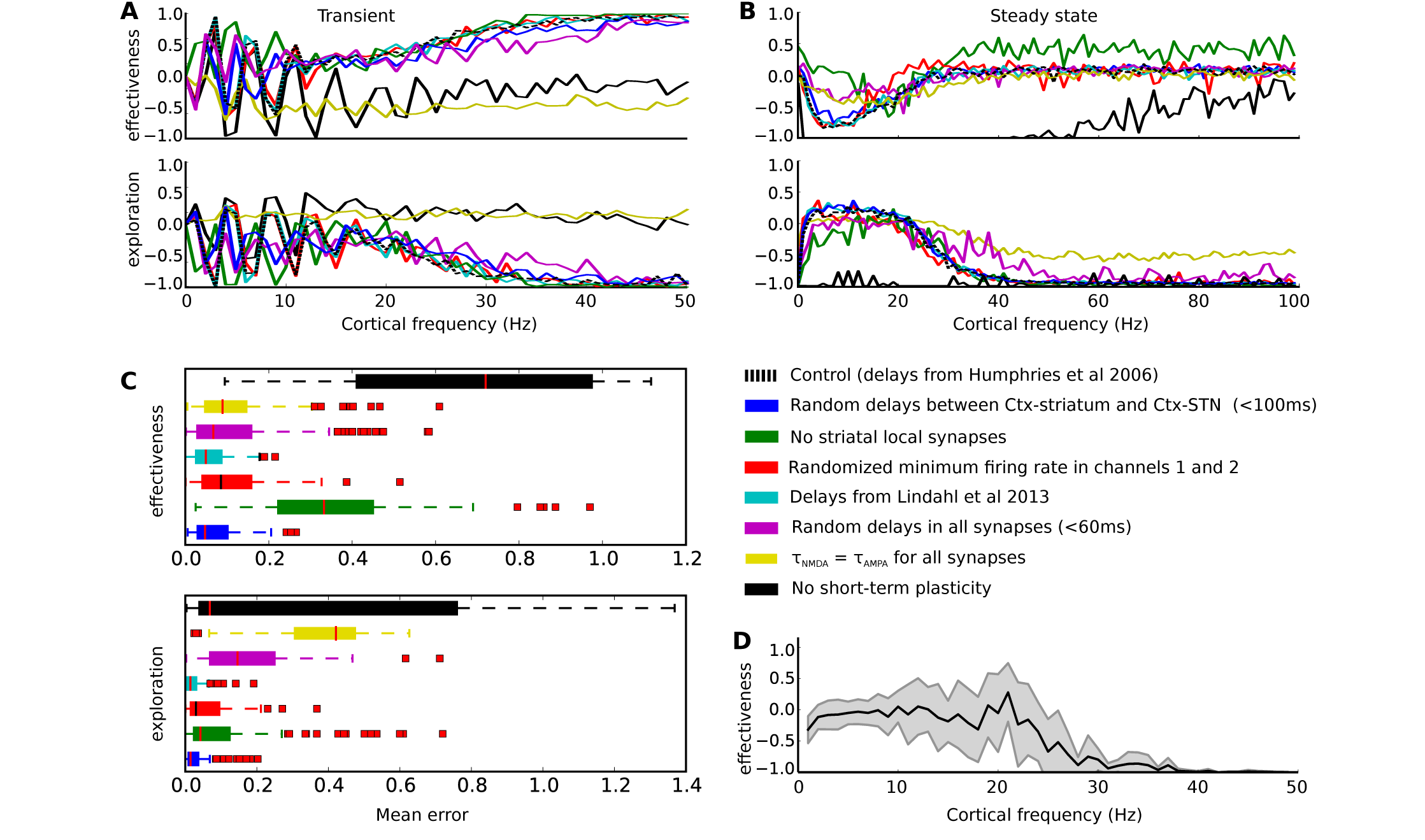
Comparison of selectivity portraits of reduced versions of the BG model. **A:** Transient effectiveness and exploration for various cortical frequencies and zero phase offset between two cortical input signals, for different versions of the BG model where a single key feature has changed. The difference between the amplitudes of these signals is 2.5 spikes/sec. **B:** The same figures but steady-state effectiveness and exploration. **C:** Box plot of the mean error between the selectivity behaviour of the default BG model and the examined reduced versions, for various initial conditions in (B). **D:** The same effectiveness figure for the minimal BG model.

The model variations that were chosen to be shown here are the ones that showed the highest differences in either effectiveness or exploration. To maintain consistency with the previous figures, we ran simulations for both sets of amplitudes *A*_1_ = 7.5, *A*_2_ = 10 and *A*_1_ = 5, *A*_2_ = 10 spikes/sec. In all cases, the feature of the model that clearly had the strongest impact on selectivity was the existence of plasticity in the chemical synapses. When plasticity was “off”, the conductance strength of the affected synapses was maintained in a static state, where the connectivity of the model was tuned to represent the baseline activity of the BG nuclei [26]. This synaptic stationarity reduced dramatically the ability of the model to make selections at any frequency, and completely impaired its ability to maintain selection for longer than 500 ms. See the black selectivity curves in Fig. 7A and B. In contrast, the lack of lateral connectivity in the striatum had a significant positive effect in steady-state selectivity, but not transiently. Finally, the selectivity of the model underwent a similar dramatic reduction with plasticity when no NMDA receptors were used in the model (*τ_NMDA_* = *τ_AMPA_*), consistent both in transient and steady state.

Interestingly, variations in conductance delays in synapses between, or within, the nuclei did not play an important role in modulating the selectivity portraits. Delays were either completely randomized, maintaining a biologically plausible range, or altered in synapses where our initial choice was based on evidence with conflicts among independent studies. For example, a computational model of the BG microcircuit presented in [30] integrated data previous studies and and concluded that the conductance delay in synapses from the STN to the GPe is on average 5 ms, for GPe-STN also 5 ms, for STN-SNr 4.5 ms, for MSN_*D*1_-SNr 7ms and for MSN_*D*2_-GPe 7 ms. In the current study, these parameters were taken from [4] where their corresponding values are 2 ms, 4 ms, 1.5 ms 4 ms and 5 ms respectively. In addition, a second example comprised changes only in the delay of the input between the cortex and STN, which represents the extra distance that information signals have to travel to arrive to the hyper-direct BG pathway. This is an important parameter of the model, since it is not yet clear what cortical areas activate the same microscopic channels in the striatum and STN. In both examples, random variations in the synaptic delays did not cause significant variations in the selectivity portraits.

Another important observation in the current comparison is the effect of the phase offset *φ* on selectivity during low-oscillations. Fig. 7A and B include curves of average selectivity over various initial conditions, but with *φ* always being fixed at zero. We chose to show these curves in order to highlight the great impact of the phase offset at low frequencies, which remained consistent among the most versions of the model. As an exemption, when no slowly-decaying synapses are used (*τ_NMDA_* = *τ_AMPA_*), this effect disappears.

Finally, Fig. 7D illustrates that the minimal version of the BG model produced a completely different behaviour. This can be attributed to a wide range of differences between the two models including the number of neurons and membrane potential dynamics. Yet, even under these simplifications, cortical oscillations at 20 Hz remained the most critical borderline in selectivity portraits that divides the frequency spectrum into two bands with antithetical behaviour (Fig. 4.Aii and Bii).

Taking everything into account, our results indicate the important hazards of oversimplification in computational modelling based on spiking neurons, since the latter do not always fall into the same level of biological abstraction.

#### The effect of the phase offset between low-frequency cortical inputs on selectivity portraits

So far, we showed that the combination of cortical frequency with the level of dopamine in the system defines how effective the BG circuitry is in discriminating incoming cortical information signals. Oscillators of various frequencies emerge constantly in the cortex, made by task-dependent coalitions of neural areas, or ensembles [20,41]. These flexible neural populations are transiently being engaged (or coupled) and disengaged (or decoupled) in a metastable manner [42], distinguishable by their different relative phases. Evidence indicates that by staying out of phase, these ensembles maintain representations of different entities in working memory [19].

Hence, it is likely that cortical groups that project to different microscopic channels in the level of the BG are phase-locked with a non-zero phase offset, which plays an important role in maintaining the identity of the potential action that is currently represented. Furthermore, since evidence points to the beta frequencies as the main range that mediates the formation of new ensembles [20], it is particularly important to assess the BG behaviour in this range.

In our simulations, we found that the phase offset *φ* between coherent cortical signals with different amplitudes can have a strong influence on the effectiveness of the BG, at certain low frequencies, while in gamma band this effect disappears (Fig. 8A). Indeed, the strongest effect was clearly located in the beta range, where the BG effectiveness was significantly enhanced when the phase of the one input signal preceded in time the phase of the second, with a small offset around 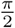.

**Fig 8.**
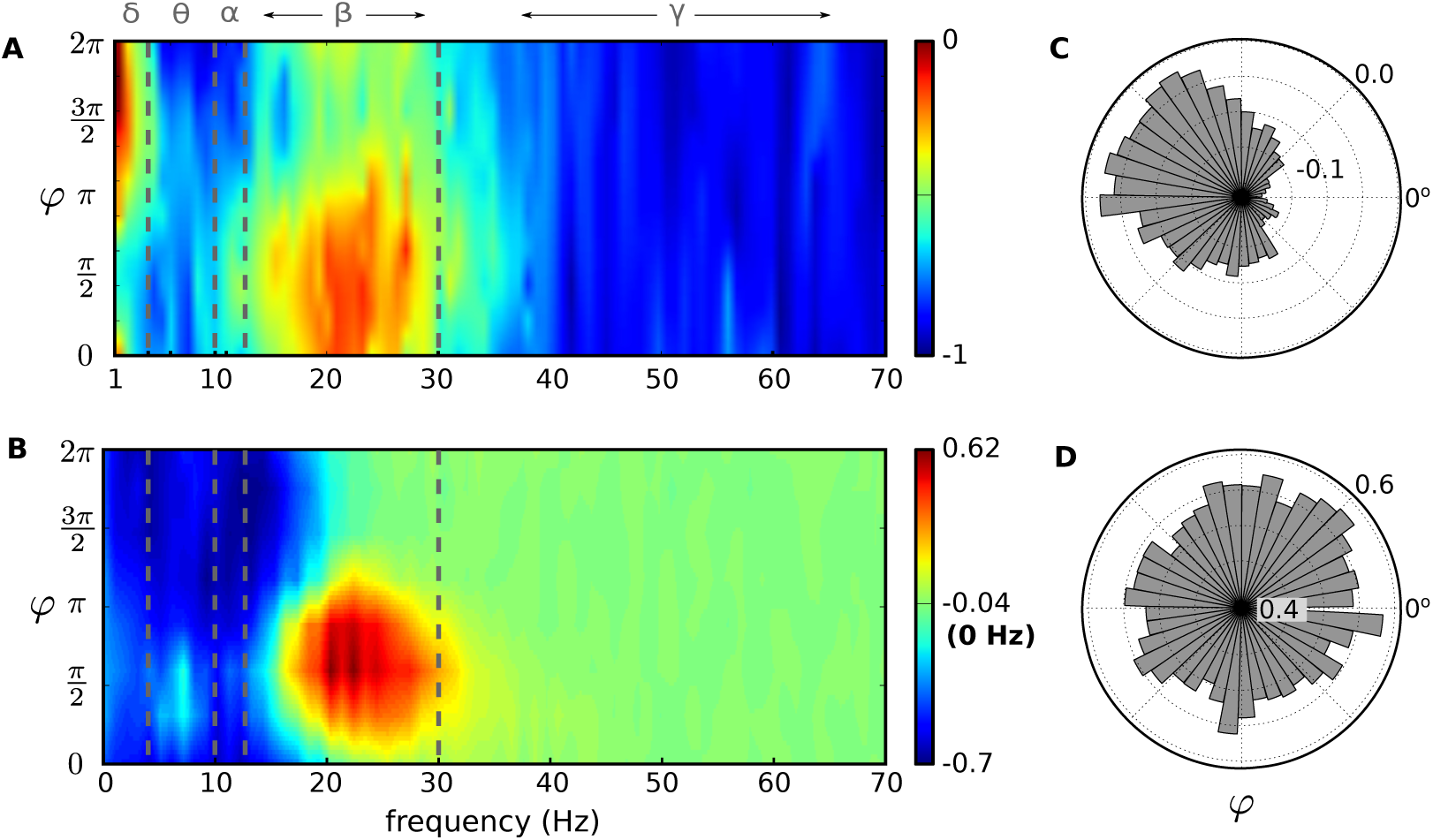
The ubiquitous effect of the phase offset *φ* at beta frequencies. **A:** BG effectiveness as a function of the frequency of two oscillatory cortical groups for *A*_2_ – *A*_1_= 2.5 spikes/sec and the phase offset *φ* between them. **B:** The same figure for the minimal model presented in Materials and Methods. **C:** BG effectiveness versus *φ* at theta frequencies. **D:** BG effectiveness versus *φ* at gamma frequencies.

Surprisingly, the sensitivity of the BG to different phase offsets during beta oscillations was largely preserved in all versions of our computational model including the minimal version. Fig. 8A illustrates this similarity which is even more prominent, since the two models produced different selectivity portraits, as a result of their numerous differences.

The relationship between phase and the BG function was investigated experimentally by [43], who showed that neural synchrony increased in the Parkinsonian BG for certain phase differences between beta oscillations in STN and GPe. Our computational model has shown that, above the alpha range, most GPe neurons that are part of a phasic microscopic BG channel remain largely silent during this phasic process [26]. Hence, the remaining GPe neurons are vulnerable to entrainment by weaker cortical inputs. As cortical beta oscillations were also shown to maintain coherence throughout the BG circuit, it is likely that the phase difference that Cagnan and her colleagues observed in this study reflected specific phase alignments of two competing cortical populations.

#### Selecting the most salient input does not require coherence between competing populations

Our results highlighted the impact of the frequency and phase of cortical ensembles that project to the BG. In order to draw conclusions regarding the phase difference between competing populations, we confined our simulations to populations of equal frequencies. However, EEG studies show that several different bands can coexist in the same or different regions of the cortex and interact with each other [44]. Hence, to explore the dynamics of BG selectivity that emerge during a combination of two stimuli with non-equal frequencies we ran another set of simulations for frequencies 0 < *f*_1_, *f*_2_ < 50 Hz and random offset *φ*. The resulting portraits are given in Fig. 9.

**Fig 9.**
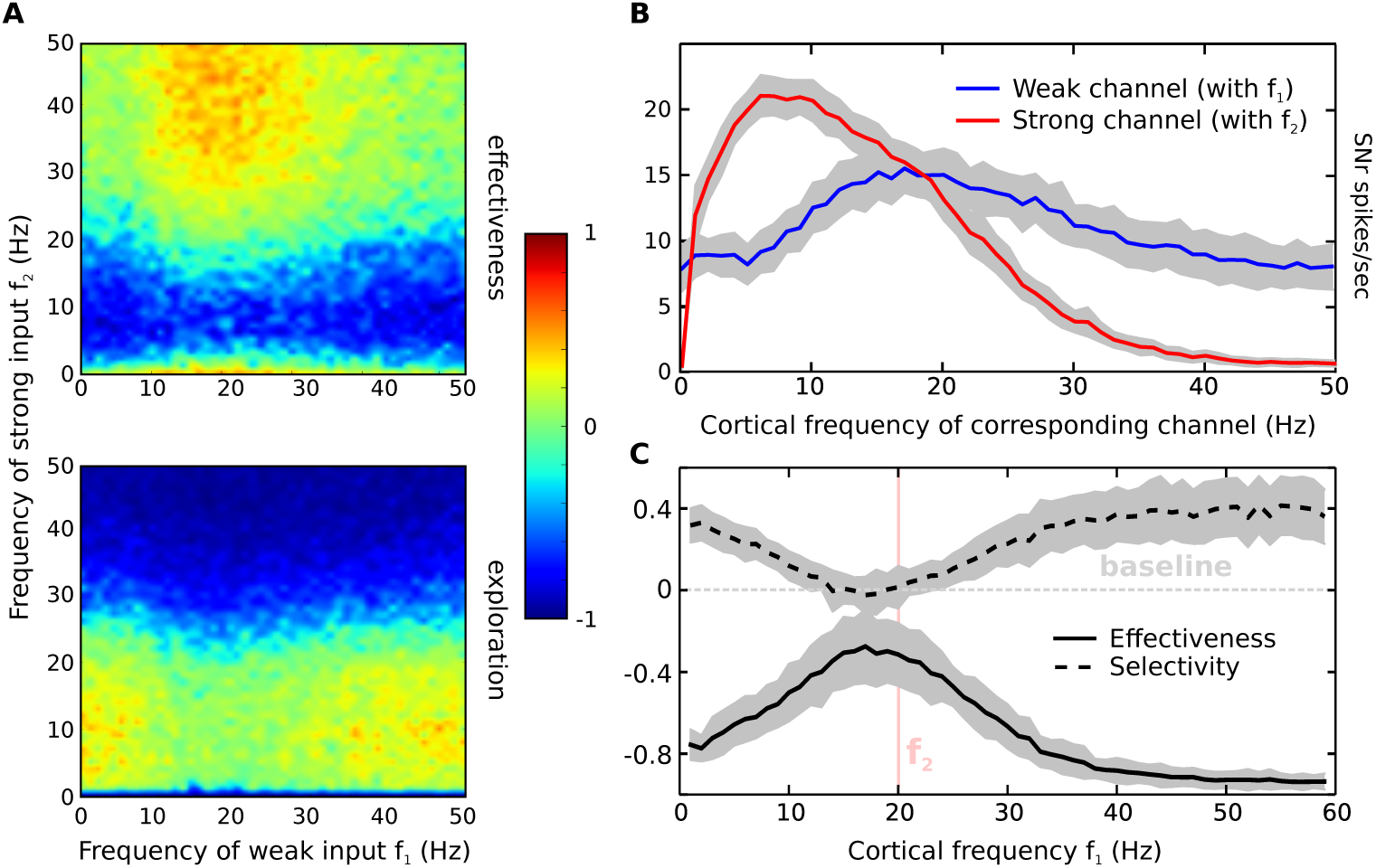
Cortical stimulation at two non-equal frequencies. **A:** Average steady-state effectiveness and exploration for all combinations of input frequencies below 50 Hz. **B:** Firing rate of the SNr in the two stimulated channels across the spectrum of the cortical frequency connected to the same channel. **C:** Average steady-state effectiveness and selectivity of the model when *f*_2_ = 20 Hz.

Despite the fact that our BG model contains various synaptic pathways that connect the two neighbouring channels, the SNr activity of each channel was immune to frequency changes in the other (Fig. 9B). Changes in effectiveness and exploration were both largely dominated by the frequency *f*_2_ of the strongest input, and across the *f*_2_ spectrum they followed a pattern similar to the portraits in Fig. 4. The oscillation of the weak channel was able to ‘bend’ this pattern only at beta frequencies, where effectiveness was enhanced.

### Behavioural predictions

#### Evidence for the existence of a long selection cycle that can be used for evidence accumulation

It is assumed by a variety of models that cognitive operations in the brain require a fixed duration [45–47], which is often referred to as a cognitive cycle. Studies have implicated the BG as the central cognitive coordinator which works in a serial manner with a cycle of 50ms [46,48].

In order to investigate the contribution of our model to this hypothesis, we simulated a two choice task experiment, following the methodology in [31]. The BG model was stimulated with tonic input of 3 spikes/sec for 1 second in order to converge to an “inactive” steady-state where no selection is being made (Fig. 10). Then, a ramping increase, which lasted for 50 ms, changed the cortical firing rate of the one channel to 10 spikes/sec (*channel 2*). A second neighbouring channel received the same increase for the first 25 ms of the ramping time, but it decayed back to its tonic firing rate after another 25 ms (*channel 1*). The cortical activity in these two channels represented the urgency for two competing actions, which in the latter less-salient case was suppressed after some initial evidence accumulation.

**Fig 10.**
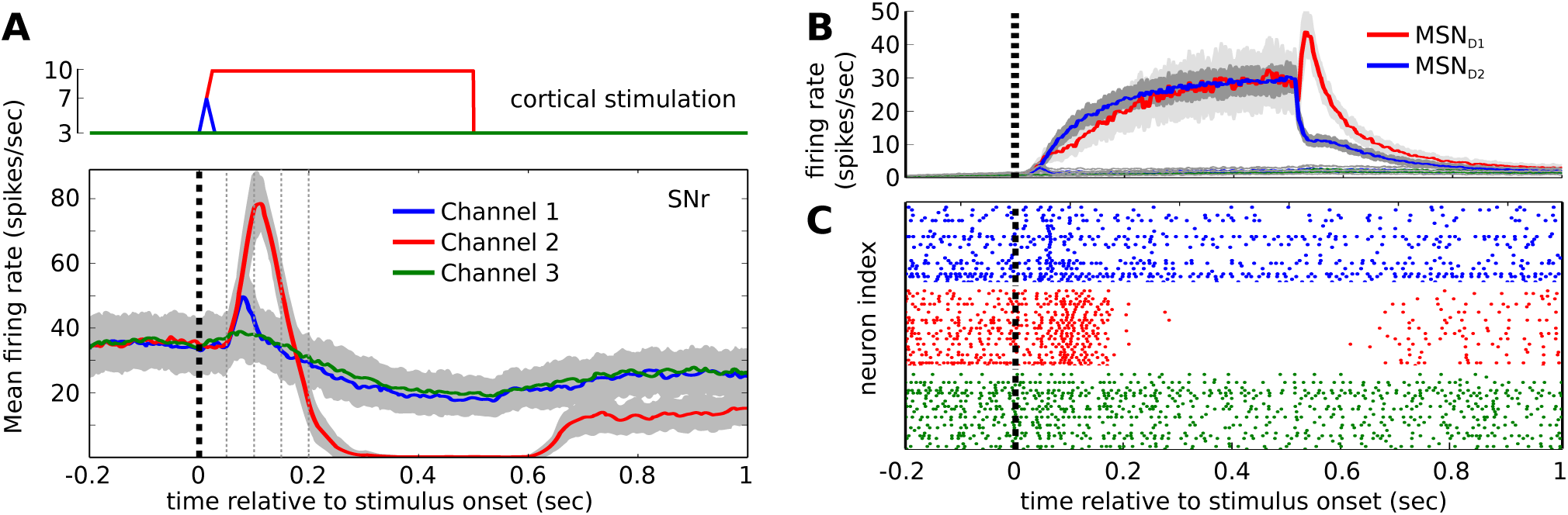
BG output during a two-choice task. **A:** Average firing rate of the 3 SNr microscopic channels and the corresponding cortical firing that caused this behaviour. **B:** The same for the two types of MSN neurons. **C:** Raster plot of the SNr during one of the runs in A and B.

Although the model of the striatum that has been used in this study is based on the neuron equations presented in [31], we observed a consistent bimodal selectivity pattern that was different from the results in this study. Since our model does not include any feedback connections from other nuclei to the striatum, this difference can be only attributed to the asymmetric inhibition between *MSN_D1_* and *MSN_D2_* neurons which was examined in [26], but it was not taken into account in [31].

The response of the SNr, the BG output nucleus, to this stimulation comprised a sequence of events. The first event occurred after 50 ms from the presentation of the stimuli in channels 1 and 2. Initially, a rapid increase in SNr firing rate was evoked, which was proportional to the intensity of the stimulus in each channel. This increase maximized after approximately 50 more ms, to be followed by a complete shut down of the selected channel, for the rest of the duration that the stimulus was presented.

The timing of this sequence of events was very similar to the experiment in Fig. 5 where the salience of the second action remained fixed during stimulation. This effect was shown to be robust and not influenced by the oscillatory patterns of the cortical input, therefore indicating the existence of a series of cognitive operations that take place during the selection process.

As shown in Fig. 10, after approximately 75 ms from the stimulation onset, channel 1 ceased to influence the outcome of the selection. But was channel 2 already selected at this particular point of time? Since the SNr does not stop its inhibitory effect to the thalamus before 200ms have passed, it is possible that a large portion of this time is used to accumulate information related to this selection. The fact that extra inhibition is provided to the phasic channels in the thalamus via the SNr, agrees well to this hypothesis.

To investigate these questions, as well as the tolerance of the time interval that is required for a successful selection, a new set of experiments was conducted. After the initial 50 ms ramp period, channel 2 received a fixed (non-oscillatory) input that had a random duration between 1 and 750 ms, while channel 1 received the same ramped input as before. In all runs, the distinctiveness *D_j_* of the three simulated channels was recorded across time, in order to see when the maximum point of effective selection can be reached in each case. The results are presented in Fig. 11.

**Fig 11.**
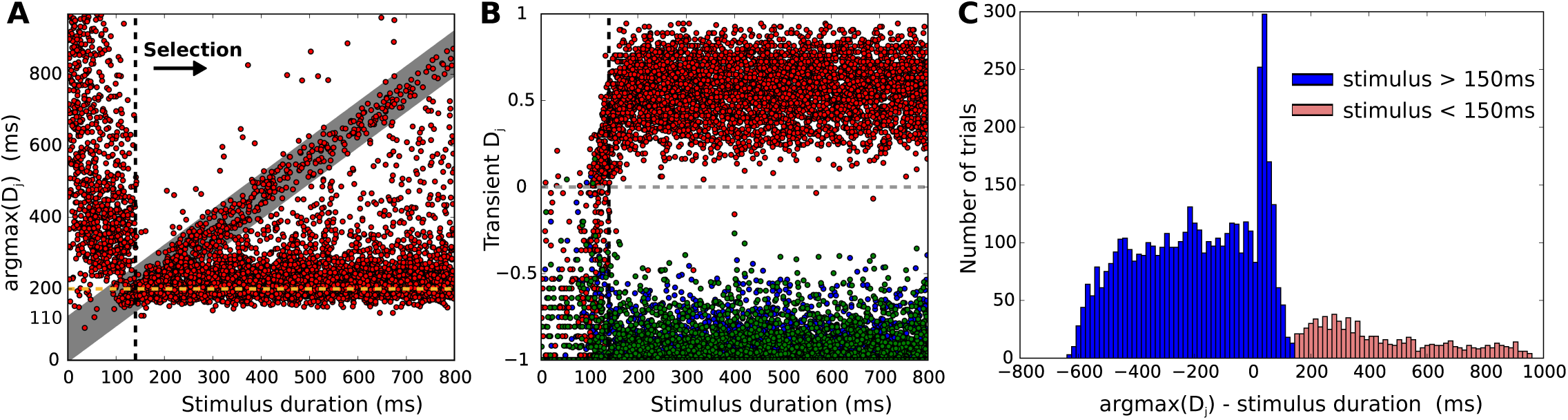
BG response for stimulus of varying duration. **A:** The point in time where BG effectiveness maximizes for a range of stimulus durations. **B:** Distinctiveness of the tree SNr channels for the same range. **C:** Histogram that shows when the maximum effectiveness was recorded with respect to the point in time that stimulation stopped.

Interestingly, we found that the BG model can discriminate between phasically and tonically-active channels only when the stimulus is presented for more than 140 ms (black dashed line in Fig. 11A and B). Longer stimuli are adequate to initiate this selection process, which normally lasts approximately 200 ms (yellow dashed line in Fig. 11A). Therefore, the inhibition of the selected channel in the level of the SNr is always preceded by excess excitation when a successful selection is performed.

This long interval, during which some information channels in the thalamus are completely shut by SNr inhibition (Fig.10A), could allow a mental deliberation process to be performed in the cortex, while the latter remains partly isolated from the environment. If during this process a channel looses its salience, as in the case of channel 1 in Fig. 10, its SNr activity will return to a neutral state, thus avoiding any interference with the final selection. Additionally, if no channel is able to maintain strong cortical activity, the process of selection will be cancelled and the excess excitation in the SNr will again prevent the inhibition of the thalamus. These features make the observed behaviour a good candidate mechanism for serial action selection.

Furthermore, the model exhibited a strong rebound effect after phasic cortical stimulation stopped. Within the range of 0 to 110 ms after stimulation, which is represented by a gray zone in Fig. 11A, the SNr inhibition of the most salient channel remained suspended. In fact, after approximately 50 ms the distinctiveness of the stimulated channel peaked again, as the neighbouring microscopic channels regained activity (Fig. 11C). This post-stimulation increase in selectivity was strongly facilitated by the rebound behaviour of the direct pathway, via excitation of MSN neurons in the striatum. As shown in Fig. 10B, the MSN_*d*1_ sub-population exhibits a sharp increase in their firing rate, which is inversely proportional to the rate of MSN_*d*2_ neurons of the same channel. Since MSNs do not evoke rebound spikes when stimulated in vitro [49], this activity can only be due to the fast decrease of local inhibition and the asymmetric connectivity between the two types of MSN neurons.

Although striatal lateral inhibition is crucial for the observed pattern of prolonged selectivity, it is not the only mechanism that causes rebound responses. Fig. 12 illustrates the response of the BG model for stimulus of various duration, when the simulated microscopic channels are connected with weak local striatal connections, to cover the possibility that these channels are physically located far from each other in the level of the striatum.

**Fig 12.**
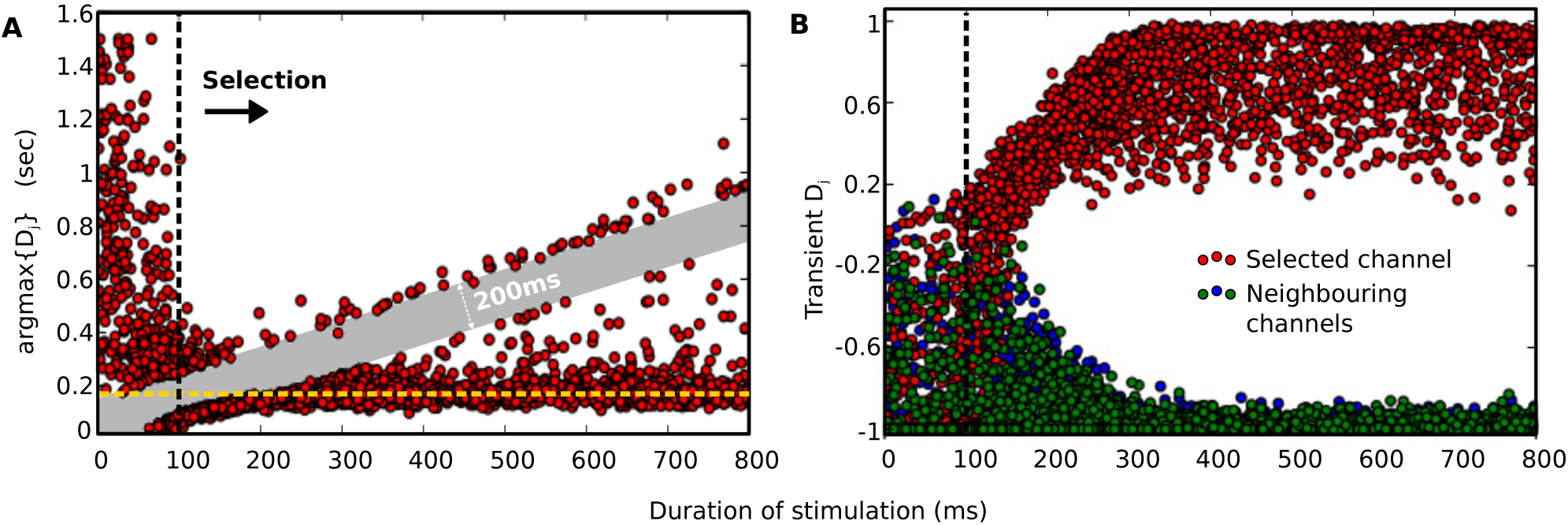
BG response for stimulus of varying duration in non-neighbouring channels. **A,B:** The same as in Fig. 11 in a variation of the BG model for weak striatal lateral inhibition that represents longer distance between channels.

In this case, the model required shorter presentation of the stimulus in order for a selection to be performed (around 100 ms). However, after stimulation stoped, it underwent a refractory period of approximately 200 ms (gray area in Fig. 12A), after which effectiveness peaked again. The fact that the duration of this period matches the initial time that the model needs to execute a selection provides additional indications of a selection cycle, which can be initiated after major changes to the input that the model receives. Although the existence of a cycle is consistent with all data presented in this section, a refractory period was not observable when strong competition took place within the striatum. A possible reason is that as the BG become more effective in distinguishing between channels, they are able to maintain a ready-to-select state, rather than initiating a new cycle, since the winning channel is already inhibiting the surrounding areas.

Finally, in order to conclude that the series of events which led to selection in our experiments constitute a cycle, the ability of the model to maintain effectiveness sequentially needs to be established. The steady-state selectivity portraits presented in Fig. 4 demonstrate that a single selection can not be maintained for many cycles of a duration longer than 70 ms (lower than beta frequencies), even if it is significantly more salient than an alternative choice.

Hence, to test if such a selection cycle can be repeated indefinitely, we ran an experiment where the three channels of the BG are stimulated sequentially for a fixed period *T* per single cycle. We found that the BG was able to distinguish the most salient channel via excitation in the SNr when *T* > 30ms. However, the second phase of the selection process, where a selection is executed via inhibition of the corresponding SNr channel, could not be achieved when *T* < 140ms. A cycle of 200 ms was able to maintain inhibition to the SNr for approximately 50 ms. These results match with the model’s behaviour in Fig. 12 and verify that the above selection process can be sequenced.

### Cognitive architectures

Although recent cognitive models are consistent with various experimental studies, a strict definition of the timing of a cognitive cycle is a challenging task. For this reason, cognitive architectures do not currently agree on a common timing model that accounts for perception, cognition and action selection [47]. As mentioned before, the BG are considered to be a fundamental element of this triad [48], which makes the model that is described in this study a useful source of information for this quest. Even with an ideal design, however, a BG model is inadequate for capturing the timing of a complete cognitive cycle, since a significantly wider range of brain structures are typically involved in this process. Alternatively, the current model can be used to impose a number of biological restrictions and to establish whether the current cognitive models can be supported by the BG dynamics (Fig. 13A).

**Fig 13.**
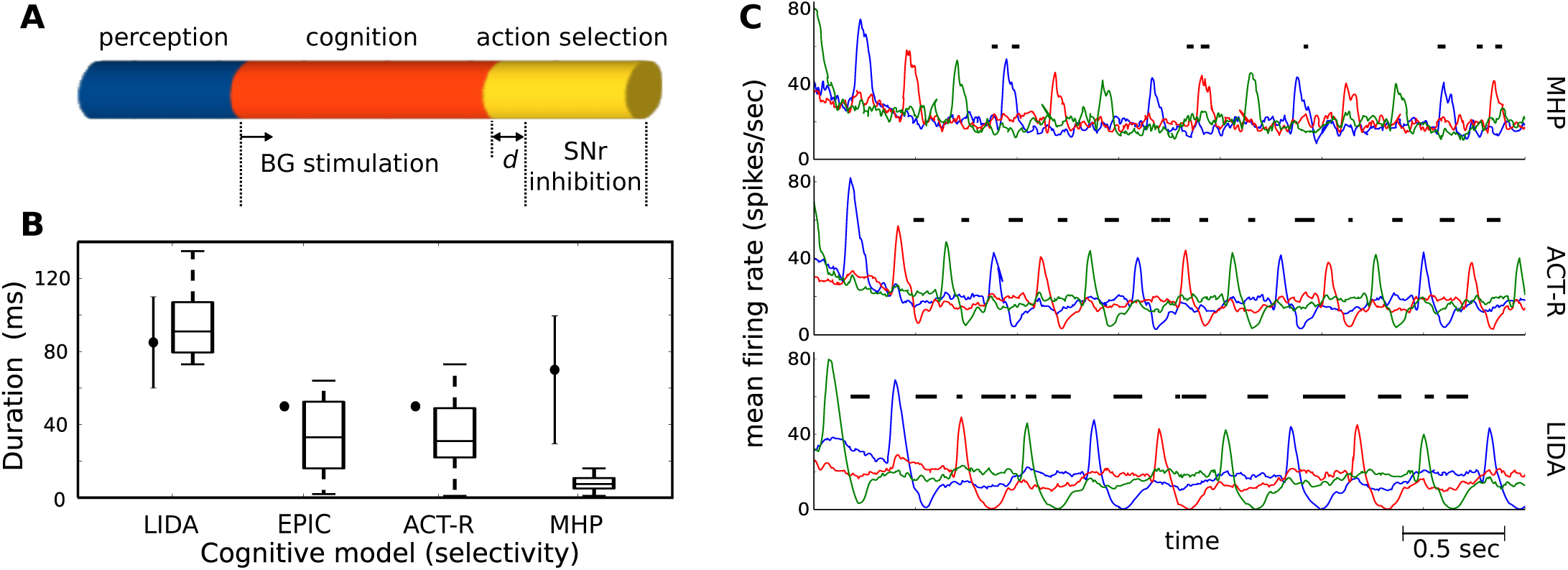
Timing of action selection in popular cognitive architectures. **A:** The three fundamental processes in a cognitive system, mapped to the activity of the BG model. The distance *d* ≥ 0 to show that selection has not occurred before the onset of this process. **B:** Distribution of durations in each cycle that selectivity is above the threshold 0.22. The black dots represent expected duration while the box plots show the results of our simulations. **C:** SNr firing rates when the BG is stimulated sequentially, with timing that matches three characteristic examples of cognitive architectures. The black lines represent areas where BG selectivity is above the threshold. The three coloured curves represent different microscopic channels.

An important restriction implied by our simulations is that a cognitive cycle should be at least 200 ms, which is the time it takes for the BG to complete a selection cycle spontaneously, measured from the onset of cortical stimulation. Although it is not clear to what extend the perception process can overlap with the activation of the cortical areas that project to the BG directly, it is safe to assume that there is a minimal overlap, given the hierarchical structure of information processing in the cortex [50]. Hence, if no parallelism between different cycles is assumed, our model suggests that a biologically plausible borderline range for the period of a cognitive cycle is from 200 ms to 200 ms plus the time duration required for perception. This restriction contradicts the majority of the currently proposed cognitive architectures, whose timing assumptions can be found in [47] and are summarised below.

One of the most popular models examined here is called Adaptive Control of Thought-Rational model (ACT-R). Originally introduced by [51], ACT-R is a modular and symbolic system which proposes that human knowledge comprises declarative memory chunks, and procedural rules. The brain is thought to be coordinated based on these rules via a central production unit, which was later associated with the function of the BG [48]. ACT-R assumes that the time of human perception is approximately 85 ms, while 100 more milliseconds are required for the rest cognitive operations before selection. These intervals can be further broken down into 50 ms cycles of production rules, which correspond to information travelling through the BG-thalamo-cortical loop. Finally, since action selection is realized as a production rule cycle, it also lasts for 50 ms and, as a result, the time that remains for the BG to process input and execute a selection is 150 ms.

While this duration is shorter than the current predictions, the 50 ms cycle of ACT-R is, to some extend, consistent with our model’s behaviour. Fig. 10 illustrates that all significant events in SNr activity that led to selection occurred in 50 ms intervals. Although the means of selection in the BG is typically hypothesized to be inhibition, excess SNr excitation discriminated the most salient microscopic channel prior to inhibition. This behaviour contradicts previous BG models and indicates that a selection is initially made in the first 100 ms, while other necessary operations take place until the selection is executed at approximately 200 ms from the stimulus onset.

A second model examined here is called Executive Process/Integrative Control (EPIC) [52]. The architecture and core assumptions of this model are very similar to ACT-R. The main difference in timing between these two models can be found in perception, which in EPIC is thought to last for 50 ms. Hence, the same conflict between ACT-R and our results applies also to this model.

Another influential approach was proposed by [53] and is called Learning Intelligent Distribution Agent (LIDA). LIDA is based on Bernard Baars’ model of consciousness named global workspace theory, according to which, conscious cognitive content is broadcasted to all active brain processes via a globally available workspace (see the theatre metaphor in [54]). LIDA assumes that perception takes 80-100 ms, the rest (unconscious) processing before action selection takes approximately 100-200 ms, while the action selection sub-process takes 60-110ms [47]. The predicted timing of a cognitive cycle proposed here falls within the limits of this theory although, on average, the duration of non-perception processes is 35 ms longer than predicted.

Finally, the Model Human Processor (MHP), proposed by [55], is based on the same division of the mind as perceptual, cognitive and motor subsystems (or processors), which are partially coupled and have different durations. A number of studies have concluded that the cycle time for the perceptual processor in young adults is on average 100 ms with a range between 50 – 200 ms depending on the task, for the cognitive processor 70 ms with a range between 25-170 ms and for the motor processor 70 ms with a range between 30-100 ms. For a review on this topic, as well as the time changes in older adults see [56]. Again, most of the range of estimated time for cognition and action selection is inconsistent with our results, which ideally require at least 140 ms for the stimulus to be projected to the BG and 60 additional milliseconds for action selection.

One issue that was not taken into account in this analysis is a potential parallelism of different cognitive cycles. Although this is a common limitation among the majority of the above models, it is known that the brain can process different tasks using some form of parallelism. Experiments with two different choice tasks performed on a single trial, have highlighted that the processing required for these tasks can overlap, but the reaction time of the second task will depend on the duration of the overlap [57]. This phenomenon, known as the psychological refractory period, is often attributed to the existence of a central bottleneck in the flow of information, that allows parallelism in perception and action execution but not during the time when the action is being selected [58]. As shown previously in this section, our BG model could support such parallel operations which can reduce the period of a cycle down to 140 ms, the time required for stimulus presentation. Thus, given the complex dynamics of decision making which are highlighted with this paradigm, further analysis is required to assess the plausibility of the above cognitive models.

In an additional experiment, the three channels of the BG were stimulated sequentially as before, for a cycle *T* equal to the proposed period of each cognitive model. Stimulation was applied only in the time interval between perception and action selection, to keep consistency between the models. The response of the model was timed, in order to investigate whether it will maintain a selection for the duration assumed by each model (Fig. 13A). To measure selectivity, we used the metric *S* which is defined in (4) and the model was considered to be actively selecting when *S* > 0.22. A comparison between original estimations of selection and the resulting durations that the SNr selected channel remained inhibited can be found in Fig. 13B.

The plausibility of LIDA was enhanced as the BG model was able to achieve the highest levels of selectivity in all trials, under these time restrictions. The timing of LIDA was also a close match, with almost all trials resulting in durations within the estimated range. Fig. 13C illustrates the average firing rate of the BG output during the trials and allows the comparison between models. Furthermore, the time restrictions of ACT-R and EPIC also allowed the BG model to exceed the threshold of selectivity. However, EPIC was a better match temporally, causing inhibition to the selected channel for 41 ± 28 ms, and also achieved higher selectivity scores.

On the other hand, when the BG model was stimulated with the temporal restrictions of MHP, action selection did not occur at all (Fig. 13C). This indicates that despite the fact that this cognitive model is able to fit to experimental data with a high degree of accuracy [56], its underlying theory may require adjustments to be biologically consistent.

### Low-frequency oscillations facilitate the resolution of ambiguity

Fig. 14 illustrates in more detail the impact of different cortical frequencies for any amplitude difference between stimuli, which represents all possible stages of a single selection. The one extreme case of *A*_1_ = *A*_2_ = 10 spikes/sec corresponds to two equally silent inputs, while the combination of *A*_1_ = 3 and *A*_2_ = 10 spikes/sec reflects the case that only one input has remained above the baseline. According to the selectivity portraits and this figure, at the beginning of a selection and when the correct choice is ambiguous, the BG are able to start exploring the most salient input only when the cortex does not oscillate at low frequencies, or during the combination of high beta and dopamine.

**Fig 14.**
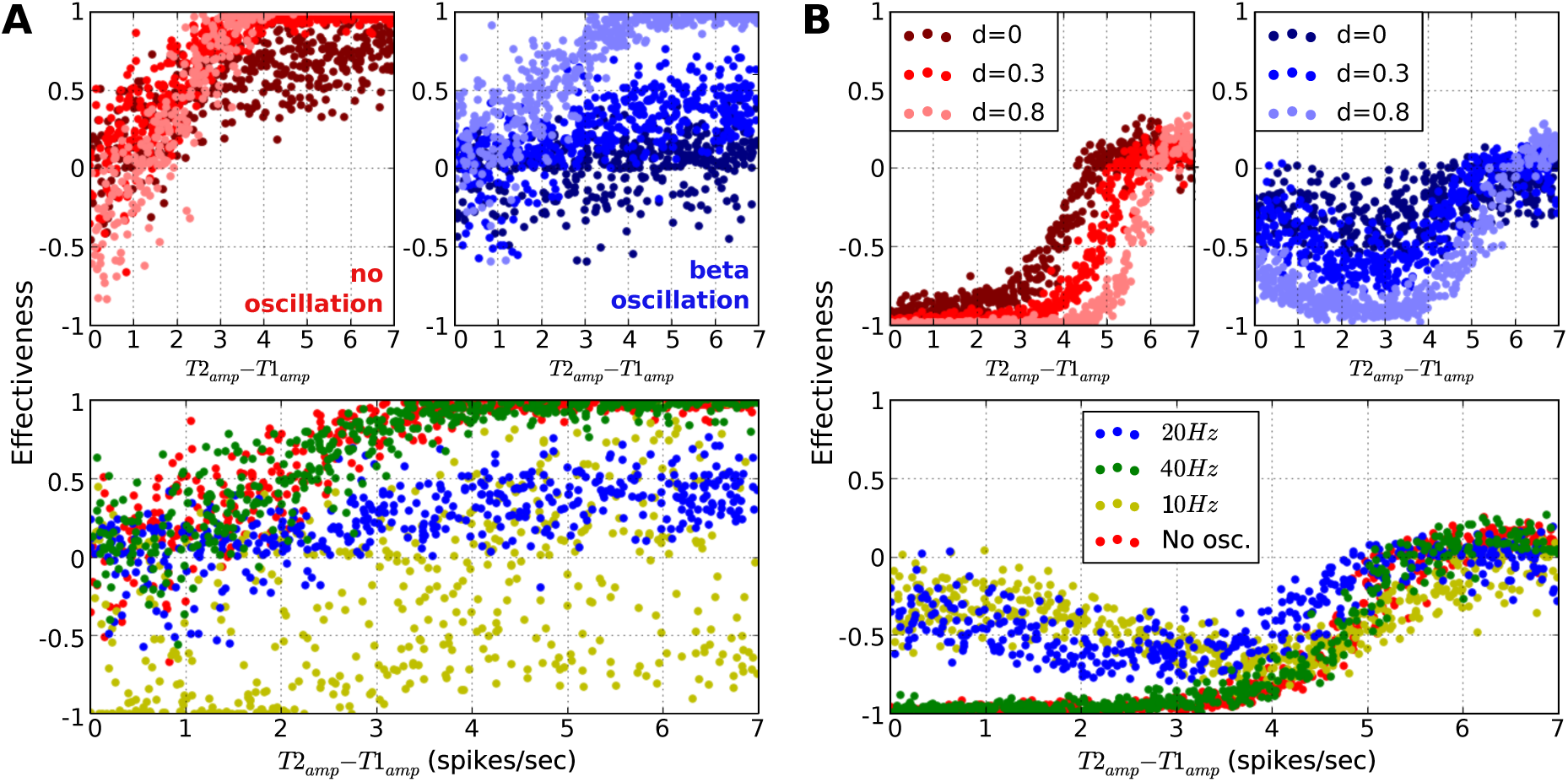
Ambiguity effects on BG selectivity. **A:** Transient effectiveness when two competing inputs have various amplitude difference, and *A*_2_ > *A*_1_. Different sets of data points represent reference cortical frequencies and dopamine levels, and are illustrated in different colours. **B:** The same as A for steady-state effectiveness.

On the other hand, if a selection task requires a longer interaction with the BG, low oscillations can maintain effectiveness near the baseline (Fig. 14B), possibly securing extra time for evidence accumulation. Also, an increased level of dopamine in this case has the opposite effect. Interestingly, the system is unable to achieve a high effectiveness score after the initial transient period, even in the case of a clear winner. This indicates that either decision making in this case is achieved on another brain region, or that long interactions for single cognitive tasks are simply not possible. If the former hypothesis is true, low-oscillatory input to the BG could facilitate selection by maintaining a neutral state among phasically-active inputs. Finally, it is worth noting that the gamma band had the same impact with no oscillations in all simulated scenarios.

All in all, the non-linear behaviour of the BG effectiveness that is illustrated in this figure, during the transition from ambiguity to certainty, shows the complexity of this circuit, when stimulated with low-frequency oscillations. However, the predictive power of our model is limited by the lack of other important brain regions, which makes difficult to draw conclusions that reflect complete behaviours.

## Discussion

### The gear box metaphor

This resulting selectivity portraits of our model constitute an interesting finding as they indicate that the cortex is in fact the structure that determines whether decision making will be performed, while the BG just execute the selected decision policy. Taking this into account, we present a novel hypothesis that views the BG as the “*gearbox*” of action selection in the brain (Fig. 15), a mechanism that provides various modes of signal selection on demand. Following this metaphor, the level of dopamine can be likened to the “*control pedals*” of action selection that either stop or initiate a decision (see selectivity portraits in Fig. 4). In the same context, the frequency of cortical oscillations acts as a “*gear lever*”, that instead of controlling the type and direction of thrust that the throttle provides to an automobile, it dictates the degree to which dopamine can trigger a decision, as well as what type of decision this would be (either exploit, stop or explore).

**Fig 15.**
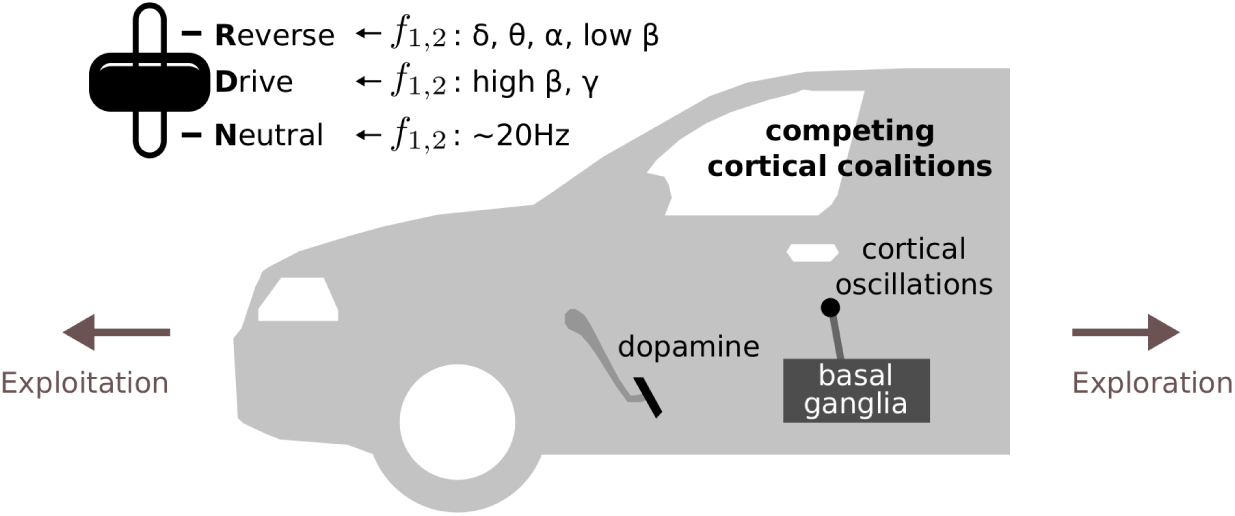
The gearbox analogy.

This framework provides justification to a number of experimental findings. Cortical beta is found to increase when a postural challenge is anticipated [59]. Since cortical beta, at around 20 Hz, can bring the BG in an *neutral* state that cancels out effectiveness and exploration, it can be viewed as a frequency that causes a temporary deactivation of the action selection system when the current action needs to be maintained. In addition, the transient effects of selectivity agree well with the duration of increases in extracellular dopamine after SNc discharges *in vivo* recordings of the rat striatum in [60,61]. A single discharge increases dopamine for approximately 200 ms while an SNc burst causes an increase that lasts about 500 – 600 ms. During this interval, our model can almost always select the most salient action (see results) and it can be significantly benefited by an increase in dopamine. However, the same selection can be maintained after this interval only if the level of dopamine decreases (Fig. 4Bii).

One advantage of using this metaphor is that it highlights a number of similarities between these two highly complex and rather unrelated dynamical systems and thus provides an intuitive way of viewing the biological mechanism of action selection.

### Relation with psychophysical studies

Our model’s predictions are also consistent with a wide range of experimental studies on mental chronometry. Although reaction times (RT) of young adults in simple tasks are in the order of 190 – 220 ms [62], these reactions can be simply stimulus-driven [63] and thus, they may bypass the action selection system of the brain. In contrast, when different responses are required depending on the class of the stimulus, choice reaction times (CRT) are found to be significantly longer, on average 500 ms in two-choice tasks [38], between 390 – 470 ms when the subjects aim for high speed, between 450 – 610 ms when the aim is high accuracy [64] and at a minimum of 200 ms [36] below which, responses are random.

By subtracting the average RT that is required by an individual to perform a simple task from the CRT of a specific choice task, we can estimate the time that is spent for the cognitive processing of the choices. This is found to vary significantly across different age groups, with an average range between 200 – 400 ms and minimum at approximately 150 ms [38]. The latency of our BG model is comparable with the lower values in the range of central processing times found in this study. This is an acceptable result given that the simulated task that was performed here constitutes arguably one of the simplest possible selection scenarios, and that the pathway of voluntary actions that involves the BG may also comprise a number of regions, as reviewed in [63], that are not simulated here.

Furthermore, a range of studies links the duration of stimulus presentation with choice task performance in mammals. In [65], monkeys were presented with visual cues of varying duration, and their accuracy on a two choice task was recorded. When the stimulus was clear, their performance increased almost linearly from near-random in viewing time of 100 ms, up to 200 ms and then there was a minimal improvement. In a *GO* odor task with rats, [66] showed that performance decreases to near random if the odor sample is presented for 100 ms or less, unless the subjects were anticipating the identity of the stimuli or the time of the response.

### Alpha and theta oscillations act as a BG mechanism to reset selection and explore alternative actions

In the literature, there is cumulative evidence that strong alpha power is able to inhibit task-irrelevant regions in the cortex and thus control information flow [15,20,67]. This theory, which is known as gating by inhibition [68], proposes that strong alpha activity is caused by GABAergic interneurons, which silence neuronal firing by providing a pulsed inhibition. Although a recent MEG study provides initial evidence that links gamma peaks to alpha troughs in the temporal cortex [69], a number of important questions still remain unanswered. For example, it is not yet clear to what extend the phase-amplitude coupling that was observed in this study was a result of local GABAergic inhibition, or other brain regions, and whether this mechanism can operate in the same spatial scale that is required to inhibit complete neural ensembles.

Based on our simulations, we propose that alpha-induced inhibition of neural populations is mediated by the selection circuit of the BG. In particular, we found that alpha and theta cortical frequencies stop the selection of the strongest input completely and instead promote the selection of less salient areas. This exploratory behaviour was independent of amplitude difference between the two inputs, occurred transiently and remained active, even after a long exposure to the stimuli (see selectivity portrait in Fig. 4). In addition, the robustness of this effect to different background frequencies was established in Fig. 9D. When the most salient input was oscillating at alpha rhythms with frequency around 10 Hz, a second weak oscillatory input was always favoured, especially when its frequency was not in the beta range.

This view of cortical alpha is consistent with a number of experimental studies. [70] recently showed that effective connectivity from the cortex to the nucleus accumbens, a part of the striatum, increases during alpha oscillations, and reverses during theta. Also, [20] presented evidence where beta synchronization in the prefrontal cortex mediated the formation of neural ensembles that represented procedural rules, while alpha synchrony increased in the ensembles that represented alternative rules. This led the authors to suggest that “beta-frequency synchrony selects the relevant rule ensemble, while alpha-frequency synchrony deselects a stronger, but currently irrelevant, ensemble”.

While alpha importance has been already discussed, the role of theta is less clear. Interestingly, the period of a theta cycle (150 – 250 ms) fits well to the timing of an action selection cycle found in our results, and it is within the limits of the full cycle of the majority of the proposed cognitive models. However, in our simulations, providing strong stimulation the model for less than 140 ms did not evoke a selection unless multiple inputs were presented sequentially. Could this be an indication that cortical theta brings the BG to its extreme limit of time efficiency, below which no selection can be achieved? In behavioural experiments, theta is found to increase in the rat striatum during a decision-making task [71], while in humans, theta in STN increases during sensorimotor conflicts [72].

### Cortical frequency is a better predictor of the exploration-exploitation trade-off than dopamine

It has been suggested that tonic dopamine levels in the striatum encode the degree of which the brain selects the action with the most predicted outcome, over the exploration of an alternative less-safe choice, by modulating activation of the direct and indirect BG pathways [73]. Fast manipulation of the trade-off between exploration and exploitation is critical for behavioural flexibility in dynamic environments [74]. This hypothesis is supported by evidence with genetically modified mice, where increased dopamine levels resulted in selections that were less influenced by the potential cost of each choice [75].

Here, the ratio between exploration and exploitation can be estimated via the ratio between distinctiveness of the most salient microscopic BC channel and distinctiveness of the rest active channels, that is, the ratio between effectiveness and exploration as defined in (3) and (5).

As shown in Fig. 4, we found that cortical rhythms play a more decisive role in this trade-off than the level of dopamine, although the combination of both cortical frequency and dopamine was crucial for the final selection. Whereas alpha and theta frequencies clearly promoted exploration over exploitation, unless uncertainty is very low, and the lack of them had the opposite effect, the level of dopamine could be largely viewed as an extra boost that triggers the selected action. In particular, during cortical beta oscillations of approximately 20 Hz, the system was in a critical state below which exploration was favoured over exploitation. However, at this very critical point and under high uncertainty, the level of dopamine was the decisive factor of the trade-off.

This complex interaction of dopamine with action selection justifies the lack of a widely accepted model, despite the fact that dopamine is evidently implicated in both exploration and exploitation [76]. On the other hand, cortical oscillations have also started to receive some attention on this topic. In [21], Cavanagh et al found a strong correlation between theta oscillations in frontal regions and uncertainty-driven exploration. This led the authors to the hypothesis that frontal areas of the cortex take over action selection from the BG in tasks with high uncertainty. Our results however show that the BG could potentially cope the need for exploratory behaviour, in case that frontal areas ‘request’ it because high uncertainty is detected.

